# Exposure to the RXR agonist SR11237 in early life causes disturbed skeletal morphogenesis in a rat model

**DOI:** 10.1101/774851

**Authors:** Holly Dupuis, Michael Andrew Pest, Ermina Hadzic, Thin Xuan Vo, Daniel B. Hardy, Frank Beier

## Abstract

Longitudinal bone growth occurs through endochondral ossification (EO), controlled by various signaling molecules. Retinoid X Receptor (RXR) is a nuclear receptor with important roles in cell death, development, and metabolism. However, little is known about its role in EO. In this study, the agonist SR11237 was used to evaluate RXR activation on EO.

Rats given SR11237 from post-natal day 5 to 15 were harvested for micro-computed tomography scanning and histology. In parallel, newborn CD1 mouse tibiae were cultured with increasing concentrations of SR11237 for histological and whole mount evaluation.

RXR agonist-treated rats were smaller than controls, and developed dysmorphia of the growth plate. Cells invading the calcified and dysmorphic growth plate appeared pre-hypertrophic in size and shape corresponding with P57 immunostaining. Additionally, SOX9 positive cells were found surrounding the calcified tissue. The epiphysis of SR11237 treated bones showed increased TRAP staining, and additional TUNEL staining at the osteo-chondral junction. MicroCT revealed morphological disorganization in the long bones of treated animals. Isolated mouse long bones treated with SR11237 grew significantly less than their DMSO controls.

This study demonstrates that stimulation of the RXR receptor causes irregular ossification, premature closure of the growth plate, and disrupted long bone growth in rodent models.

## Introduction

Longitudinal bone growth occurs through the process of endochondral ossification, in which a cartilaginous model first expands through the activity of chondrocytes, followed by replacement of cartilage by bone. Endochondral ossification begins with the condensation of mesenchymal cells that subsequently differentiate into chondrocytes, the primary cell type of cartilage. Chondrocytes begin expressing SRY (sex determining region Y)-box9 (SOX9) and synthesize a matrix comprised largely of type II collagen and the proteoglycan aggrecan [1]. These early cartilage cells have a distinct rounded shape, are scattered irregularly throughout the matrix, divide infrequently, and give rise to resting chondrocytes. Eventually, the resting cells begin to divide more rapidly, and enter the proliferative zone, although the driving force behind this shift in activity is not completely known [2]. While in the proliferating zone, cells appear flattened. Upon division, their daughter cells line up in columns along the longitudinal axis of the bone. The elongated shape and the longitudinal growth of endochondral bones are largely due to this spatial orientation [2], with chondrocyte proliferation as one key factor influencing longitudinal bone growth. Chondrocytes in the center of the cartilage eventually cease to proliferate and exit the cell cycle to become hypertrophic cells. Chondrocytes undergoing hypertrophy increase up to 20-fold in size [3], stop synthesizing SOX9, and begin expressing type X collagen instead of type II. Hypertrophic differentiation is driven by transcription factors such as runt-related transcription factor 2 (RUNX2) [4–6] and Myocyte enhancer factor-2c (MEF2C) [7] that is required for *Runx2* expression [1]. Hypertrophic chondrocytes release signals to attract blood vessels, osteoclasts invade the matrix to resorb the cartilage, and trabecular bone formation initiates. During this process, hypertrophic chondrocytes either undergo apoptosis, or may trans-differentiate into osteoblasts [1,8,9].

These processes are occurring in the center of the cartilage template, forming the primary ossification center. A secondary ossification center forms later at the end of the template through a similar process. The growth plate consists of the chondrocytes located between these two areas of ossification, with distinctly divided zones of resting, proliferating and hypertrophic cells [10]. Throughout this process, proliferation and hypertrophy of chondrocytes drive the longitudinal growth of the bone [11]. A complex array of hormones and signaling molecules control the process of endochondral ossification [12]. Dysregulation of any of the factors controlling growth plate function can cause defects including reduced bone length, altered mineralization, and skeletal deformities, such as seen in chondrodysplasias [1,5,10–12]. Understanding these processes is crucial to etiology of the bone diseases along with developing interventions to ameliorate any adverse outcomes.

Retinoid X Receptor (RXR) is a type 2 nuclear receptor that is part of the steroid/thyroid hormone superfamily. It has important roles in cell death, development, metabolism, and cell differentiation [13] and is expressed in almost every tissue of the body [14]. There are three different RXR proteins (alpha, beta and gamma), and each is expressed in a cell type-dependent manner [13]. RXR has the ability to form homodimers or heterodimers with other nuclear receptors in a permissive or non-permissive way [15,16]. Permissive heterodimers can be activated by agonists for either partner in the dimer and include RXR dimers with peroxisome proliferator-activated receptors (PPARs), farnesoid x receptors (FXRs), and liver x receptors (LXRs). Non-permissive dimers can only be activated by the ligand for the partner (not by RXR ligands) and include retinoic acid receptors (RARs), vitamin d receptors (VDRs), and thyroid hormone receptors (TRs). Given that RXR homodimerizes with itself and heterodimerizes with several nuclear receptors, thereby regulating downstream transcription of multiple genes, the direct influence of RXR alone on development has been difficult to elucidate. The RXR binding partner most broadly studied in relation to rodent development is RAR. Although this complex is important for many processes in development, the most profound effects of RXR-RAR heterodimers have been discovered in embryonic bone. For example, exposure to retinoic acid during embryonic development causes skeletal element deletions and truncations in the forelimbs [17]. While RXR is involved in these processes, the RXR-RAR heterodimer is primarily activated by an RAR agonist [18]. It remains to be studied how RXR homodimers would affect bone development without the influence of any other binding partner.

Although the impact of RXR itself is far-reaching, little is known about its relevance in long bone development. To determine its importance, the RXR specific pan-agonist SR11237 [19] was used in this study to identify the effects of RXR activation on rat endochondral bone development.

## Materials and Methods

### Animals

Sprague-Dawley rats were purchased from Charles River Laboratories (St. Constant, Quebec, Canada). Animals were housed in a controlled temperature (20-25°C), controlled humidity (40-60%) ventilated animal room. They were bred in-house and given free access to water and standard rat chow. Once born, pups were housed with their mother until harvest at post-natal day 16 (P16). All animal experiments were handled in accordance to the guidelines from the Canadian Council on Animal Care and were approved by the Animal Use Subcommittee at the University of Western Ontario.

### SR11237 Injections

In each trial, Sprague-Dawley rat pups were divided randomly into two groups. The first group was given an intraperitoneal injection (IP) of SR11237 (Sigma, Oakville, Ontario, Canada, #S8951) (pan-RXR specific agonist) at a concentration of 25 mg per kg body weight. The second group was injected with DMSO (dimethyl sulfoxide) (Sigma, Oakville, Ontario, Canada, #472301) vehicle. Neonates were injected IP once a day from post-natal day 5 to 15, in a similar regime as published with GLP (glucagon-like peptide) agonists in newborns [20], during a critical window of bone development in the rat [1,5,10–12]. This dose of SR11237 is similar to what has been previously used in vivo [21]. The animals were harvested on post-natal day 16. Animals were weighed at time of harvest.

### Tibia Organ Cultures

Tibiae were harvested from newborn CD1 mice and cultured for 4 days in a 37°C humidified chamber at 5% CO_2_ as described [22]. They were cultured in serum-free α-MEM (minimum essential medium eagle) media (Invitrogen, Burlington, Ontario, Canada, #12571-063) containing 0.2% bovine serum albumin (Fisher, Ottawa, Ontario, Canada, #BP1600-100), 0.25% L-glutamine (Invitrogen, Burlington, Ontario, Canada, #25030-081), 0.216mg/mL β-glycerophosphate (Sigma, Oakville, Ontario, Canada, #G9891), 0.05mg/ml ascorbic acid (Sigma, Oakville, Ontario, Canada, #A4034), and 0.4% penicillin-streptomycin (Invitrogen, Burlington, Ontario, Canada, #15140-122) as described [22]. On days 1 and 3, the media was replaced, and the bones were treated with treated with DMSO (vehicle) control or SR11237 at 3 increasing concentrations (0.1 μM, 1 μM, and 5 μM). Total bone length was determined immediately following isolation and upon experimental completion on day 4, using a Leica S6 D microscope with EC3 camera and Leica Application Suite version 3.8.0 software. Following fixation in 4% paraformaldehyde overnight at room temperature, tibiae were prepared for paraffin embedding and sectioning. The sections were stained with Safranin O / Fast Green as above.

### Micro-computed Tomography (microCT)

Following harvest of the P16 rats, the left limbs were removed, fixed overnight in 4% paraformaldehyde (Sigma, Oakville, Ontario, Canada, #P6148), and stored in 70% ethanol at 4°C until scanning. Males were scanned using the GE eXplore speCZT at a resolution of 50 micrometers / voxel and analyzed using the Microview 2.5.0-3943 software [23].

### Histology and Immunohistochemistry

Humerus, radius, tibia and femur were harvested and isolated from the right limbs of all animals. Long bones were measured, fixed at room temperature for 24 hours with 4% paraformaldehyde (Sigma, Oakville, Ontario, Canada, #P6148), and decalcified for 12 days with 5% ethylene diamine tetra-acetic acid (Sigma, Oakville, Ontario, Canada, #EDT001.5) in phosphate buffered saline. Bones were processed and embedded in paraffin wax, and sectioned by the Molecular Pathology Laboratory at Robarts Research Institute (London, ON, Canada) at a thickness of 5μm. Sections were dewaxed using xylenes, and then rehydrated by a series of graded ethanols. Following rehydration, sections were either chemically stained or immunohistochemistry was performed. Sections were stained with 1.5% Safranin O (Sigma, Oakville, Ontario, Canada, #S8884)/ 0.01% Fast green (Harleco (Millipore), Etibocoke, Ontario, Canada, #210-12) as described [24–26], Picro-sirius red (Picric acid saturated solution Sigma, Oakville, Ontario, Canada, #P6744-1GA and Sirius Red F3B Sigma, Oakville, Ontario, Canada, #36-554-8) as described [23], Tartrate-resistant acid phosphatase (TRAP) (Sigma, Oakville, Ontario, Canada, #387-A) according to manufacturer’s instructions with minor changes (60 minute Triton-X incubation), and Terminal deoxynucleotidyl transferase dUTP nick end labeling (TUNEL) (Calbiochem (Millipore), Etobicoke, Ontario, Canada, #QIA33) as instructed by the manufacturer’s manual. Immunohistochemistry was performed as described [24] with primary antibodies against PCNA (proliferating cell nuclear antigen) (Cell Signaling, Danvers, MA, USA, #2586) at a concentration of 1:5000, P57 (Santa Cruz, Dallas, Texas, USA, #sc-8298) at a concentration of 1:200, or SOX9 (R&D, Minneapolis, MN, USA, #AF3075) at a concentration of 1:300. Antigen retrieval was completed using 0.1% Triton-X in water, and slides without primary antibody were used as negative controls. All antibody incubations were performed overnight at 4°C. After washing, secondary antibodies conjugated to horseradish peroxidase (either Santa Cruz, Dallas, Texas, USA, #sc-2004 or sc-2020) were used for 1 hour at room temperature, followed by colourimetric detection by diaminobenzidine substrate (Dako, Santa Clara, CA, USA, #K3468). Following this, sections were counterstained using Methyl green (Sigma, Oakville, Ontario, Canada, #198080). A total of 6 sections per sample were used, with at least 3 independent samples (animals) for each assay. Images were captured directly with a Leica DM1000 microscope and Leica DFC295 camera or stitched together from multiple individual images using Leica Application Suite software.

### Statistical Analyses

GraphPad Prism software version 6 (GraphPad Software Inc., San Diego, CA, USA) was used to conduct all statistical analysis. Animals weights and bone lengths were analyzed using an unpaired two-tailed t-test with five independent trials. Organ culture tibial lengths were analyzed using a one-way ANOVA with five independent trials with Bonferonni’s post-test. Significance was assumed for P values less than 0.05 for all statistical tests.

## Results

### RXR activation results in reduced body weight and bone length

Newborn rats were injected with DMSO or SR11237 daily from P5 to P15. Daily treatment of vehicle or SR11237 had no effects on maternal behaviour or neonatal mortality (data not shown). At P16, animals were weighed before sacrifice. The RXR agonist-treated animals of both sexes weighed 30% less than the DMSO injected controls (Figure 1A, B). Humerus, tibia, femur and radius were dissected from the right side of all Sprague-Dawley pups and measured for length. An unpaired two-tailed t-test showed that all four bones were significantly shorter in the animals injected with SR11237 than the DMSO controls, in both males and females (Figure 1C, D).

**Figure 1:**
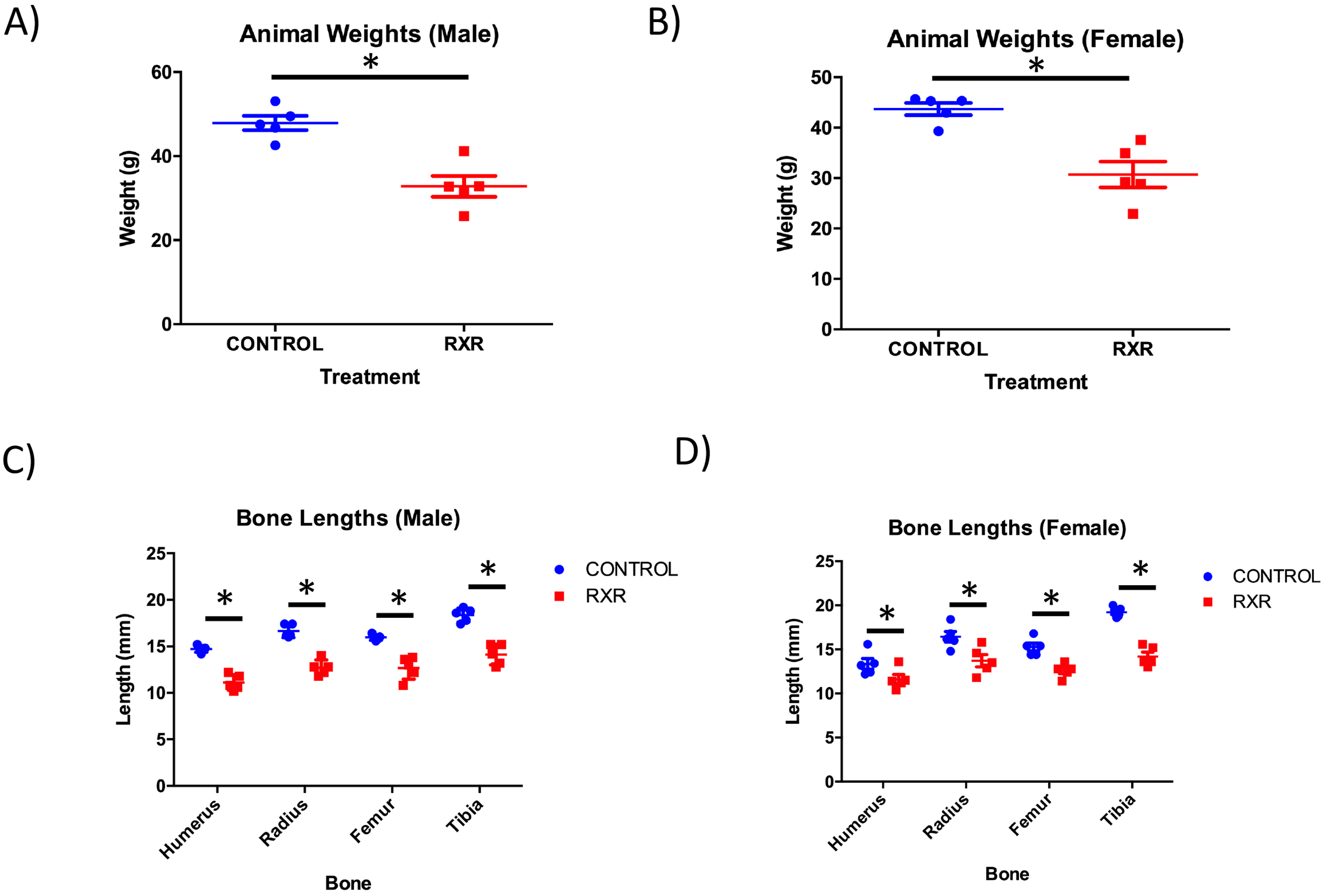
RXR Agonist Treatment Results in Reduced Weight and Bone Length in Rats. At P16, male (A) and female (B) animals were weighed before sacrifice (N=5; mean ± SEM; unpaired twotailed t-test; p<0.005). Bone lengths were measured at P16 in male (C) and female (D) rats following sacrifice. All RXR agonist-treated bones were found to be significantly shorter than control bones (N=5; mean ± SEM; unpaired two-tailed t-test; p<0.005).

### RXR Agonist SR11237 Decreases Total Length Change in Organ Cultured Mouse Tibiae

Tibiae were isolated from newborn mice and cultured for 4 days with DMSO (Control) or differing concentrations of RXR agonist SR11237. Total length of bones was measured at beginning and end of the culture period to determine longitudinal growth (Figure 2). Treatment of tibiae with 5µM SR11237 (Figure 2A) caused a statistically significant decrease in length when compared to DMSO control while lower concentrations had no effect. Safranin O/Fast Green staining of tibia sections confirmed that DMSO-treated tibiae were significantly longer than tibiae treated with 5µM SR11237 (Figure 2B).

**Figure 2:**
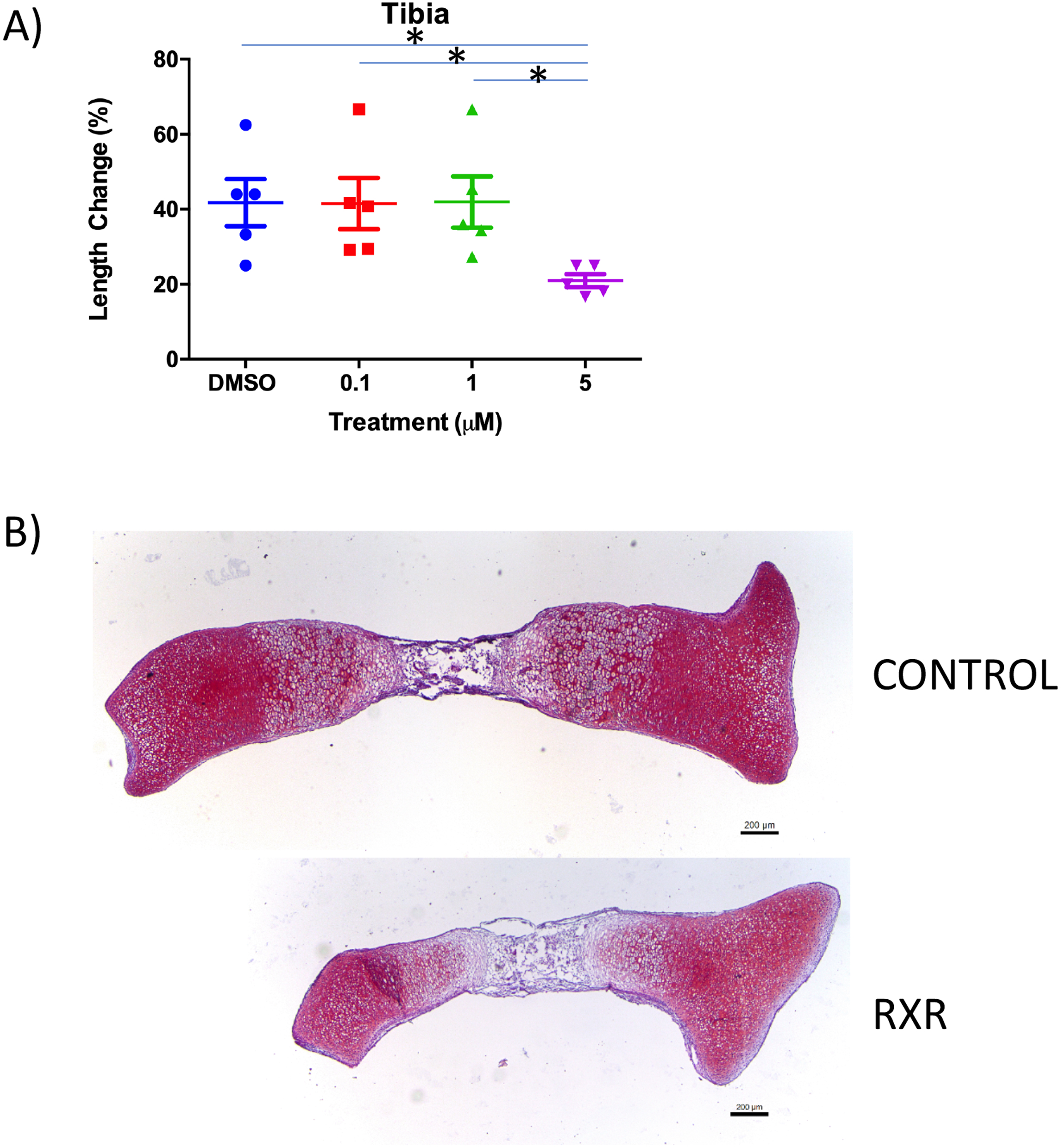
RXR Agonist SR 11237 Decreases Bone Growth in Murine Tibiae In Vitro. P0 tibiae were isolated and cultured for 4 days with DMSO and various concentrations of SR 11237. Total length of bones was measured following isolation and upon experimental completion to determine percentage of longitudinal growth. Treatment of tibia with 5µM SR 11237 causes a significant decrease in growth when compared to DMSO control (A) (N=5; mean ± SEM; one-way ANOVA with Bonferonni’s post-test; p<0.005). The DMSO (top) treated tibia is longer than the tibia treated with 5µM SR **11237 (bottom) (B).**

### MicroCT Analyses Shows Abnormal Bone Morphology in RXR agonist-treated rats

The left limbs of P16 male rats treated with RXR agonist or control were harvested for microCT analysis. The shape and size of all long bones changed substantially upon treatment with the RXR agonist. Images demonstrated a significant size reduction in the fore and hindlimbs treated with SR11237 compared to the DMSO control-treated limbs (Figure 3A, B). Regions of increased and decreased radio-opacity can be seen in all bones of the fore and hindlimbs, scapula, hands and feet upon RXR agonist treatment (Figure 3A, B, C, D, E). These scans indicated that the scapula showed less calcification in the centre of the bone, but thicker calcification along the outer rim in the animals treated with the RXR agonist (Figure 3C). Metacarpals and metatarsals of the hands and feet appeared dysmorphic, under-calcified and under-developed in treated rats (Figure 3D, E). SR11237 treatment resulted in disruption in the development of the secondary ossification centres, as well as the reduction in the size of the mineralization of the smaller bones of the hands and feet (Figure 3D, E).

**Figure 3:**
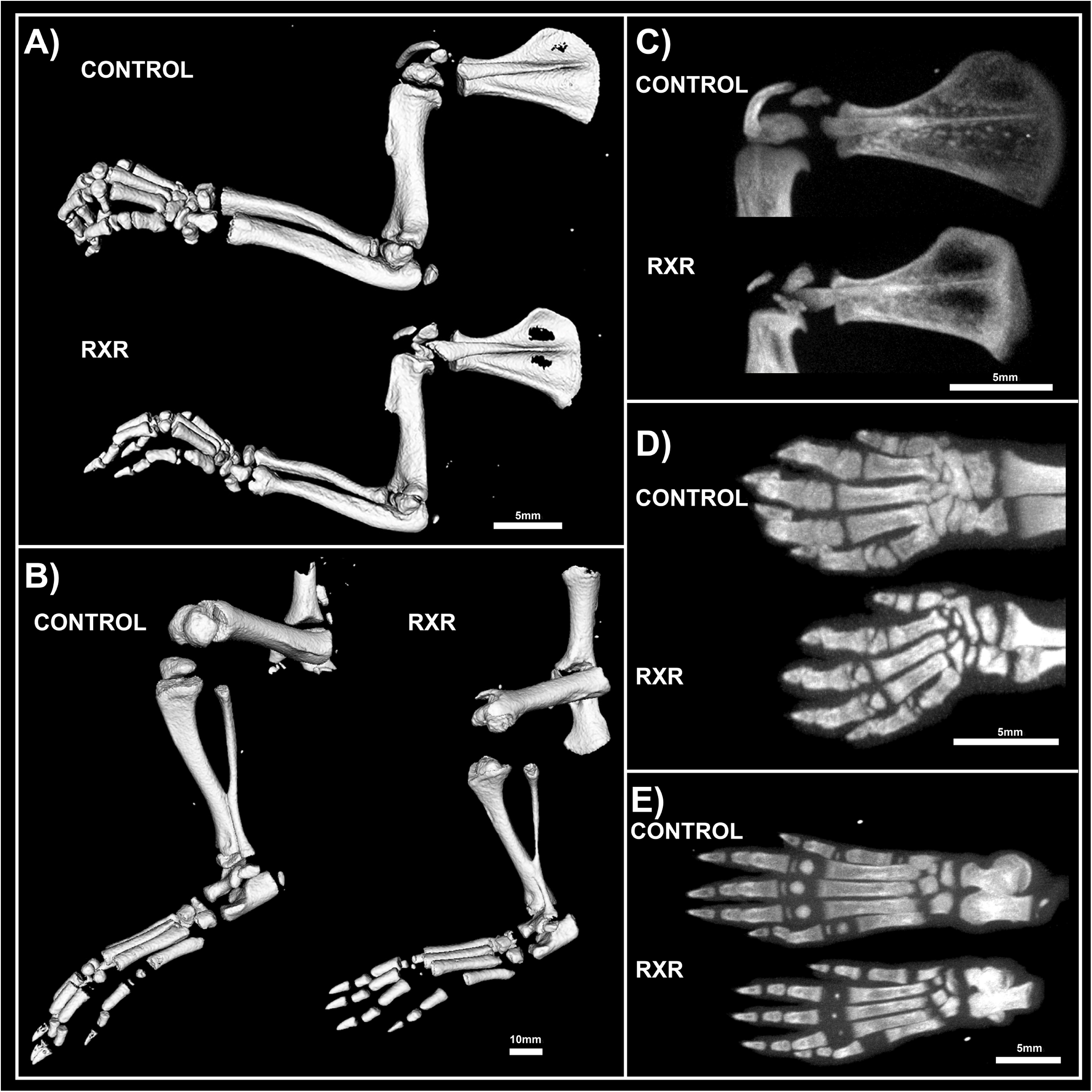
μCT Images of P16 RXR and Control Male Rats Show Abnormal Morphology. Fore and hindlimbs bones of control rats were longer and thicker than in RXR agonist-treated animals (A, B). The scapula of the RXR agonist-treated animals had increased radio-opacity in the centre of the bone, but more calcification along the outer edges when compared to the animals treated with DMSO (C). The metacarpals and metatarsals of the hands and feet appeared dysmorphic, under-calcified and under-developed in the animals treated with the RXR agonist (D, E) (N=5; 50 micrometers **/ voxel).**

### Disrupted Growth Plate Morphology in P16 Male Rat Long Bones

Paraffin sections of humerus, tibia, and femur from the P16 rats were stained with Safranin O / fast green (Figure 4). This staining highlighted greatly disturbed growth plate organization and the apparent fusion of the primary and secondary ossification centers in the RXR agonist-treated animals versus DMSO controls. RXR activation by SR11237 caused premature growth plate closure and an infiltration of ossified tissue through the central epiphysis of the bones, effectively joining together the primary and secondary ossification centres. Importantly, the tibia appeared to be more afflicted than the humerus and femur (Figure 4). Females showed a similar but less severe phenotype than males (Suppl. Figure 1).

**Figure 4:**
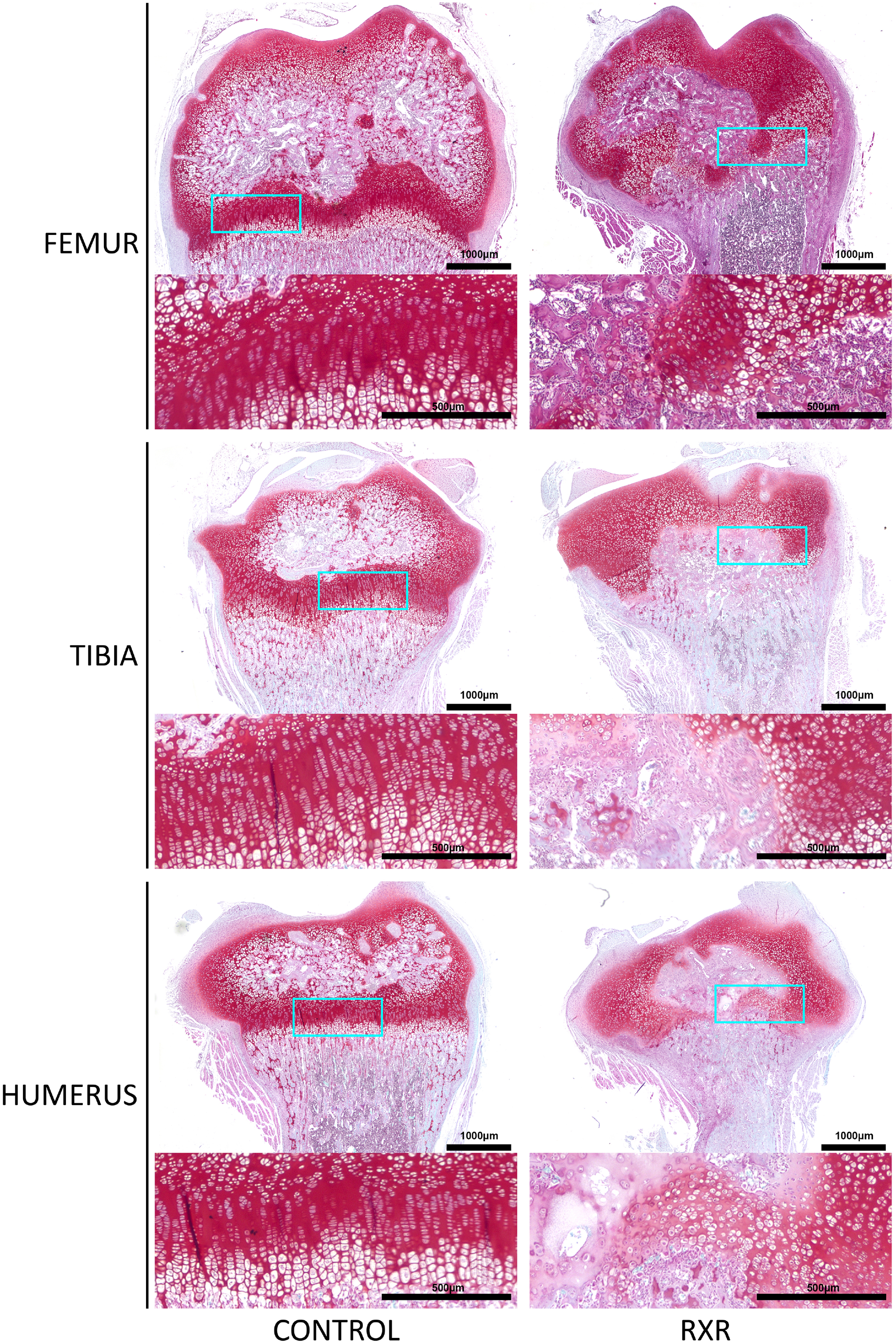
Disrupted Growth Plate Morphology in P16 Male Rat Long Bones. Safranin O / fast green staining of bone sections highlights appearance of disturbed growth plate organization and fusion of primary and secondary ossification centers in the RXR agonist-treated males (scale bar = 1000μm; inset = 500μm).

### Immunohistochemistry and Histological Staining on P16 Rat Tibial Sections

We performed histological and immunohistological analyses to further characterize the observed phenotypes. SOX9 (a marker for early and proliferating chondrocytes) expression was found in growth plate chondrocytes in control rats and in the cells surrounding the calcified tissue in SR11237-treated rats (Figure 5). The cartilage of SR11237-treated animals showed reduced numbers of chondrocytes staining positive for PCNA, a marker of cell proliferation, and P57, a marker of post-mitotic and prehypertrophic chondrocytes (Figure 5). In contrast, the central epiphysis of SR11237-treated rats was highly positive for TRAP and picro-sirius red staining, suggesting increased osteoclast activity and accelerated replacement of cartilage by bone (Figure 6). TUNEL staining appeared concentrated cell death at the osteo-chondral junction of the growth plate in both groups (Figure 6).

**Figure 5:**
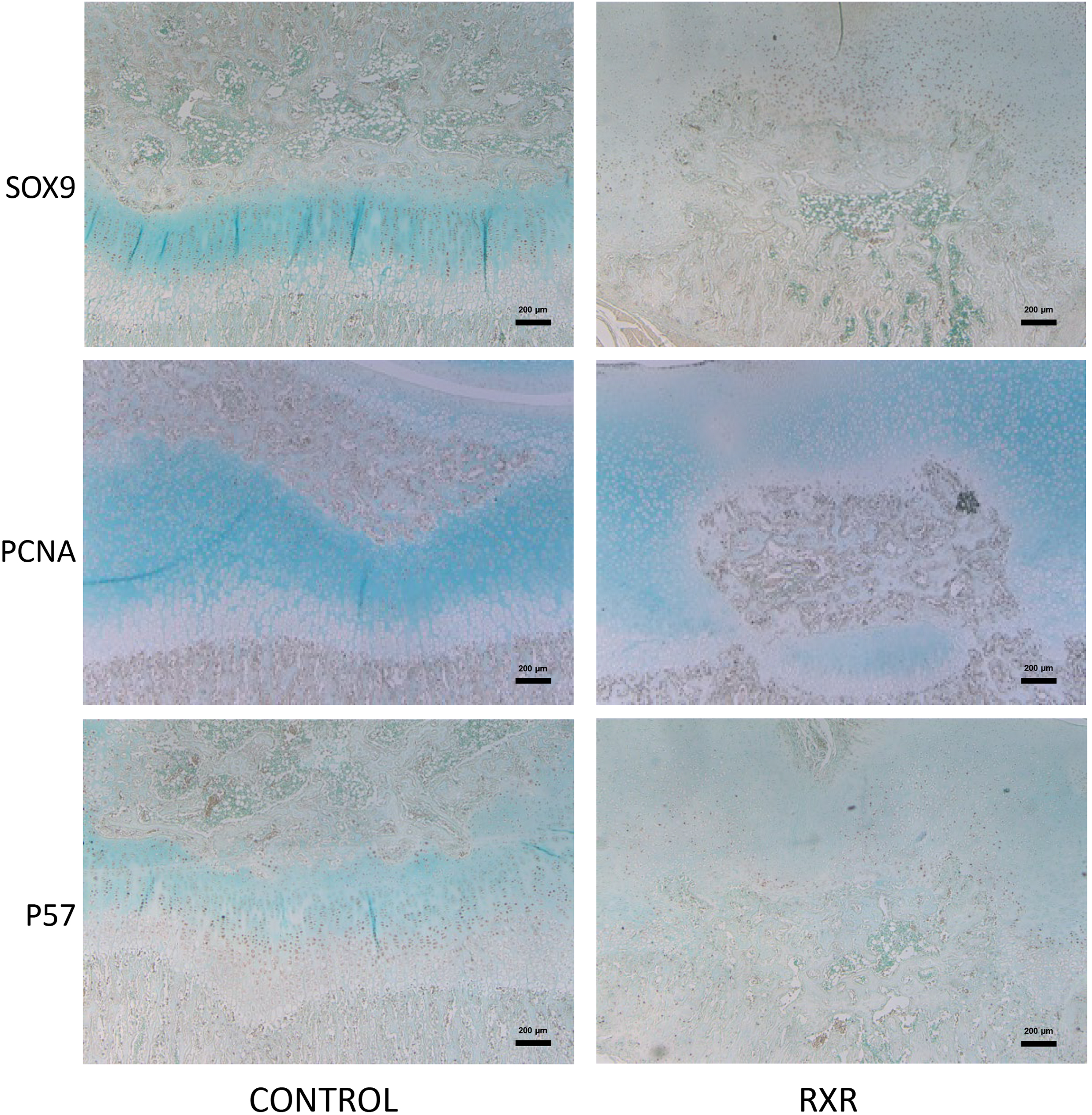
Immunohistochemistry staining on P16 Rat Tibial Sections. Sections of tibiae from rats treated with DMSO or SR 11237 were examined by immunohistochemistry for various markers. PCNA is a proliferative marker; P57 demonstrates the arrangement of terminally differentiating chondrocytes in the hypertrophic region; SOX9 shows the organization of proliferating chondrocytes (scale bar = 200μm).

**Figure 6:**
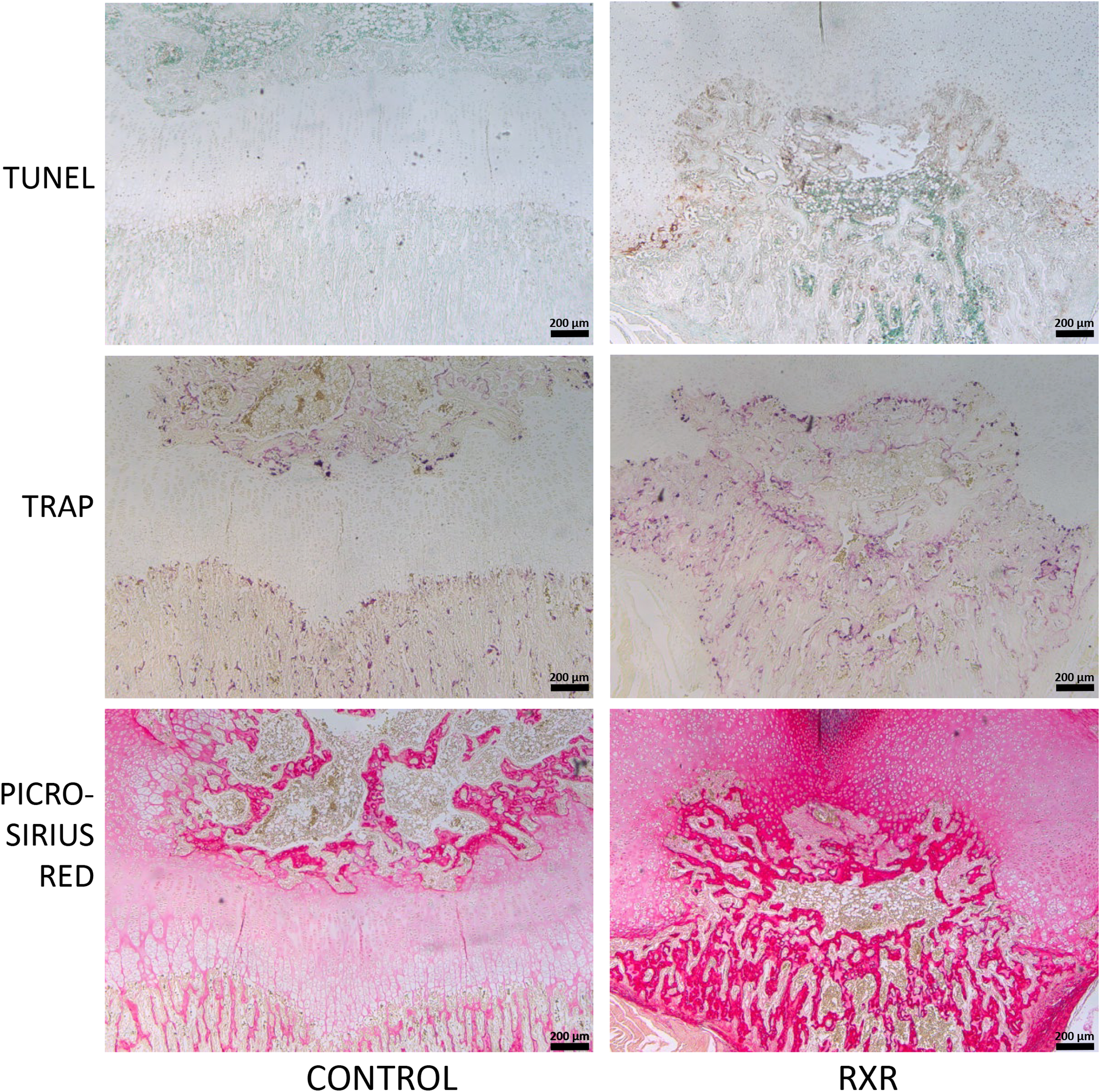
Histological staining on P16 Rat Tibial Sections. Sections of tibiae from rats treated with DMSO or SR 11237 were examined by staining for various markers. TUNEL detects cell death; active osteoclasts are stained by TRAP; Picro-sirius red stains collagen fibres (scale bar = 200μm).

## Discussion

This study examines the effects of RXR signaling on rodent endochondral ossification and growth plate biology. By activating RXR specifically in the early newborn stage, prior to and during the onset of secondary ossification, using the RXR agonist SR11237, we found adverse effects on the growth plate of rat long bones, as well as effects on body size, bone formation, and morphology of multiple skeletal elements. Previous studies had examined the effects of other RXR agonists on bone parameters but were largely focused on adult stages [27]. Similarly, mice with specific deletions of the RXRα and RXRβ genes in hematopoietic cells were studied and showed defects in osteoclast function, but were studied at later stages only [28].

The RXR agonist-treated animals of both sexes were significantly smaller (30%) than their littermate DMSO injected siblings. This size difference was present until sacrifice at P16. Similarly, all four long bones dissected (tibia, femur, humerus and radius) were significantly shorter in the animals injected with SR11237 compared to DMSO controls. At present it is unclear whether the effects on body weight are due to systemic and metabolic effects or secondary to reduced bone growth. However, the reduced growth of tibiae cultured with the high concentration of the RXR agonist suggests that at least a portion of the growth retardation observed is due to direct effects on growth plate chondrocytes.

MicroCT imaging demonstrated that the fore and hindlimbs of the SR11237-injected animals were thinner and appeared somewhat osteopenic. Skeletal staining with Alcian Blue and Alizarin Red revealed similar findings (data not shown). The metacarpals and metatarsals of the hands and feet appeared dysmorphic, undercalcified, and underdeveloped. Although both sexes appeared to be afflicted, males appeared more impaired than females (data not shown).

Safranin O/ fast green staining of long bone sections highlighted disturbed growth plate organization and the disruption of the primary and secondary ossification centers in the RXR agonist-treated animals versus the DMSO controls. Cells surrounding the disrupted growth plate and ossification centres were largely pre-hypertrophic in size and shape. Again, males appeared to be more affected than females (data not shown). Importantly, the tibia appeared to be more afflicted than the humerus and femur. In all staining and immunohistochemical experiments, sections from both male and female rats (data not shown) were used. However, in every instance, males appeared to be more afflicted than females.

P57, SOX9, and PCNA immunohistochemistry was performed on the tibia, humerus and femur sections. All three markers were present in control animals in the expected patterns, but staining patterns in the animals treated with SR11237 further demonstrated the extreme disorganization of the growth plate. Upon RXR activation, PCNA, P57, and SOX9 were found in the cells surrounding the central ossified section in no particular spatial order, suggesting that the overall organized structure of the zones of the growth plate was lost, and chondrocytes of various maturation stages are intermingled. The reduced number of PCNA-positive chondrocytes upon SR11237 administration suggests that RXR activation drives the cells out of the cell cycle, which could contribute to reduced bone growth and the closure of the growth plate.

Although it was difficult to tell if the SR11237-injected animals were more afflicted than their DMSO controls, the pattern of TUNEL staining appeared to be moderately different. Generally, the highest amount of cell death in a typically developing growth plate occurs in the late hypertrophic zone, where some chondrocytes are undergoing apoptosis and osteoclasts are invading [29]. However, in the SR11237-treated animals, TUNEL-positive cells appeared most consistently around the infiltration of ossified tissue through the growth plate, seemingly in no particular order, again demonstrating the intermingling of multiple chondrocyte types in the same space. However, there did appear to be an increase in cell death at the osteo-chondral junction, which is consistent with the normal transition as some hypertrophic cells undergo apoptosis. The highly localized pattern of TUNEL staining also suggests that the severe phenotypes observed are not due to general cellular toxicity induced by the RXR agonist.

In the DMSO control animal growth plate, TRAP staining occurs only at the site where the hypertrophic cartilage transitions to bone, and around the area of secondary ossification. In the RXR agonist-treated animals, TRAP staining is found throughout the central epiphysis of the bone and connects the primary and secondary ossification centers completely. The staining appeared very intense and prevalent throughout, suggesting that osteoclast activation or recruitment is one of the main responses to RXR activation.

Picro-sirius red stains collagen fibers and is particularly strong in bone tissue. Control animals showed picro-sirius staining concentrated at the primary and secondary ossification centers, but only weak staining in the area of the growth plate. Upon RXR activation, the area and intensity of staining appears greatly increased, further supporting our finding of increased bone formation and highlighting the fusion of primary and secondary ossification centers.

Future work will need to address whether the effects outlined in this study are due to RXR action itself or involves any of its heterodimeric partners. Potential candidates include the vitamin D receptor (VDR) since it stimulates the expression of RANKL (receptor activator of nuclear factor kappa-B ligand) in cartilage, resulting in activation of osteoclasts at the osteochondral junction [30]. In addition, deletion of the gene for RXRγ worsens the growth plate phenotype of VDR KO mice, providing additional evidence for a shared function of these receptors in cartilage [31].

Similarly, retinoic acid (potentially through retinoic acid receptors / RARs) stimulates cartilage to bone turnover in vitro [32], resembling some of the phenotypes we observed here. However, since both VDR and RAR are non-permissive partners of RXR, it seems less likely that they mediate these effects of a specific RXR agonist. Mouse knockout studies also suggest reduced osteoclast activity in mice with deletions of the genes encoding several other partners of RXR, including PPARγ, LXRs and thyroid hormone receptor alpha (reviewed in [33]); thus, activation of the respective heterodimers by our RXR agonists could contribute to the observed effects. However, it should be noted that in parallel experiments administration of the LXR agonist GW3695 did not induce any obvious bone growth effects (data not shown). Overall it seems quite plausible that the complex effects observed in our study could be a combination of activation of RXR homodimers and several heterodimeric pairs, potentially differently in the different cell types involved (chondrocytes, osteoblasts, osteoclasts and maybe more).

In conclusion, our studies demonstrated that increased activation of RXR via a selective agonist (SR11237) in skeletally immature rats caused irregular ossification, premature closure of the growth plate, and reduced bone growth during early postnatal development. Consequently, prolonged RXR signaling, especially in early life, may result in long term effects on long bone and joint morphology, and lead to additional pathology (such as osteoarthritis) through joint mal-alignment, disruption of normal gait, or direct effects on articular chondrocytes.

## Acknowledgements

FB is the Canada Research Chair in Musculoskeletal Research; research in his lab is supported by operating grants from the Canadian Institutes of Health Research (Application 332438) and The Arthritis Society (SOG-15-293). DBH was supported by a Natural Sciences and Engineering Research Council of Canada Discovery Grant (RGPIN-04090).

## Abbreviations

EO: endochondral ossification
RXR: retinoid x receptor
SOX9: sex determining region-box 9
RUNX2: runt-related transcription factor 2
MEF2C: myocyte enhancer factor–2c
PPAR: peroxisome proliferator-activated receptor
FXR: farnesoid x receptor
LXR: liver x receptor
RAR: retinoic acid receptor
VDR: vitamin d receptor
TR: thyroid receptor
IP: intraperitoneal
DMSO: dimethyl sulfoxide
GLP: glucagon-like peptide
MEM: minimum essential medium eagle
TRAP: tartrate resistant acid phosphatase
TUNEL: terminal deoxynucleotidyl transferase dUTP nick end labelling
PCNA: proliferating cell nuclear antigen
RANKL: receptor activator of nuclear factor of kappa-B ligand

## References

1. Kozhemyakina, E.; Lassar, A.B.; Zelzer, E. A pathway to bone: signaling molecules and transcription factors involved in chondrocyte development and maturation. Development 2015, 142, 817–31.

2. Abad, V.; Meyers, J.L.; Weise, M.; Gafni, R.I.; Barnes, K.M.; Nilsson, O.; Bacher, J.D.; Baron, J. The role of the resting zone in growth plate chondrogenesis. Endocrinology 2002, 143, 1851–7.

3. Cooper, K.L.; Oh, S.; Sung, Y.; Dasari, R.R.; Kirschner, M.W.; Tabin, C.J. Multiple phases of chondrocyte enlargement underlie differences in skeletal proportions. Nature 2013, 495, 375–8.

4. Takeda, S.; Bonnamy, J.P.; Owen, M.J.; Ducy, P.; Karsenty, G. Continuous expression of Cbfa1 in nonhypertrophic chondrocytes uncovers its ability to induce hypertrophic chondrocyte differentiation and partially rescues Cbfa1-deficient mice. Genes Dev. 2001, 15, 467–81.

5. Ueta, C.; Iwamoto, M.; Kanatani, N.; Yoshida, C.; Liu, Y.; Enomoto-Iwamoto, M.; Ohmori, T.; Enomoto, H.; Nakata, K.; Takada, K.; et al. Skeletal malformations caused by overexpression of Cbfa1 or its dominant negative form in chondrocytes. J. Cell Biol. 2001, 153, 87–100.

6. Yoshida, C.A.; Yamamoto, H.; Fujita, T.; Furuichi, T.; Ito, K.; Inoue, K.; Yamana, K.; Zanma, A.; Takada, K.; Ito, Y.; et al. Runx2 and Runx3 are essential for chondrocyte maturation, and Runx2 regulates limb growth through induction of Indian hedgehog. Genes Dev. 2004, 18, 952–63.

7. Arnold, M.A.; Kim, Y.; Czubryt, M.P.; Phan, D.; McAnally, J.; Qi, X.; Shelton, J.M.; Richardson, J.A.; Bassel-Duby, R.; Olson, E.N. MEF2C transcription factor controls chondrocyte hypertrophy and bone development. Dev. Cell 2007, 12, 377–89.

8. Pest, M.A.; Beier, F. Developmental biology: Is there such a thing as a cartilage-specific knockout mouse? Nat. Rev. Rheumatol. 2014, 10, 702–4.

9. Yang, L.; Tsang, K.Y.; Tang, H.C.; Chan, D.; Cheah, K.S.E. Hypertrophic chondrocytes can become osteoblasts and osteocytes in endochondral bone formation. Proc. Natl. Acad. Sci. 2014, 111, 12097–12102.

10. Yeung Tsang, K.; Wa Tsang, S.; Chan, D.; Cheah, K.S.E. The chondrocytic journey in endochondral bone growth and skeletal dysplasia. Birth Defects Res. C. Embryo Today 2014, 102, 52–73.

11. Sun, M.M.-G.; Beier, F. Chondrocyte hypertrophy in skeletal development, growth, and disease. Birth Defects Res. C. Embryo Today 2014, 102, 74–82.

12. Lui, J.C.; Nilsson, O.; Baron, J. Recent research on the growth plate: Recent insights into the regulation of the growth plate. J. Mol. Endocrinol. 2014, 53, T1–9.

13. Dawson, M.I.; Xia, Z. The retinoid X receptors and their ligands. Biochim. Biophys. Acta 2012, 1821, 21–56.

14. Pérez, E.; Bourguet, W.; Gronemeyer, H.; de Lera, A.R. Modulation of RXR function through ligand design. Biochim. Biophys. Acta 2012, 1821, 57–69.

15. Germain, P.; Chambon, P.; Eichele, G.; Evans, R.M.; Lazar, M.A.; Leid, M.; De Lera, A.R.; Lotan, R.; Mangelsdorf, D.J.; Gronemeyer, H. International Union of Pharmacology. LXIII. Retinoid X Receptors. Pharmacol. Rev. 2006, 58, 760–772.

16. Kojetin, D.J.; Matta-Camacho, E.; Hughes, T.S.; Srinivasan, S.; Nwachukwu, J.C.; Cavett, V.; Nowak, J.; Chalmers, M.J.; Marciano, D.P.; Kamenecka, T.M.; et al. Structural mechanism for signal transduction in RXR nuclear receptor heterodimers. Nat. Commun. 2015, 6, 8013.

17. Sucov, H.M.; Izpisúa-Belmonte, J.C.; Gañan, Y.; Evans, R.M. Mouse embryos lacking RXR alpha are resistant to retinoic-acid-induced limb defects. Development 1995, 121, 3997–4003.

18. Kochhar, D.M. Limb development in mouse embryos. I. Analysis of teratogenic effects of retinoic acid. Teratology 1973, 7, 289–95.

19. Benoit, G.; Altucci, L.; Flexor, M.; Ruchaud, S.; Lillehaug, J.; Raffelsberger, W.; Gronemeyer, H.; Lanotte, M. RAR-independent RXR signaling induces t(15;17) leukemia cell maturation. EMBO J. 1999, 18, 7011–8.

20. Stoffers, D.A.; Desai, B.M.; DeLeon, D.D.; Simmons, R.A. Neonatal exendin-4 prevents the development of diabetes in the intrauterine growth retarded rat. Diabetes 2003, 52, 734–40.

21. Standeven, A.M.; Escobar, M.; Beard, R.L.; Yuan, Y.D.; Chandraratna, R.A. Mitogenic effect of retinoid X receptor agonists in rat liver. Biochem. Pharmacol. 1997, 54, 517–24.

22. Agoston, H.; Khan, S.; James, C.G.; Gillespie, J.R.; Serra, R.; Stanton, L.-A.; Beier, F. C-type natriuretic peptide regulates endochondral bone growth through p38 MAP kinase-dependent and -independent pathways. BMC Dev. Biol. 2007, 7, 18.

23. Pest, M.A.; Russell, B.A.; Zhang, Y.-W.; Jeong, J.-W.; Beier, F. Disturbed cartilage and joint homeostasis resulting from a loss of mitogen-inducible gene 6 in a mouse model of joint dysfunction. Arthritis Rheumatol. (Hoboken, N.J.) 2014, 66, 2816–27.

24. Gillespie, J.R.; Ulici, V.; Dupuis, H.; Higgs, A.; Dimattia, A.; Patel, S.; Woodgett, J.R.; Beier, F. Deletion of glycogen synthase kinase-3β in cartilage results in up-regulation of glycogen synthase kinase-3α protein expression. Endocrinology 2011, 152, 1755–66.

25. Solomon, L.A.; Li, J.R.; Bérubé, N.G.; Beier, F. Loss of ATRX in chondrocytes has minimal effects on skeletal development. PLoS One 2009, 4, e7106.

26. Ulici, V.; Hoenselaar, K.D.; Agoston, H.; McErlain, D.D.; Umoh, J.; Chakrabarti, S.; Holdsworth, D.W.; Beier, F. The role of Akt1 in terminal stages of endochondral bone formation: angiogenesis and ossification. Bone 2009, 45, 1133–45.

27. Nowak, B.; Matuszewska, A.; Filipiak, J.; Nikodem, A.; Merwid-Lad, A.; Piesniewska, M.; Fereniec-Golebiewska, L.; Kwiatkowska, J.; Szelag, A. The influence of bexarotene, a selective agonist of the retinoid receptor X (RXR), and tazarotene, a selective agonist of the retinoid acid receptor (RAR), on bone metabolism in rats. Adv. Med. Sci. 2015, 61, 85–89.

28. Menéndez-Gutiérrez, M.P.; Rőszer, T.; Fuentes, L.; Núñez, V.; Escolano, A.; Redondo, J.M.; De Clerck, N.; Metzger, D.; Valledor, A.F.; Ricote, M. Retinoid X receptors orchestrate osteoclast differentiation and postnatal bone remodeling. J. Clin. Invest. 2015, 125, 809–23.

29. Kronenberg, H.M. Developmental regulation of the growth plate. Nature 2003, 423, 332–336.

30. Masuyama, R.; Stockmans, I.; Torrekens, S.; Van Looveren, R.; Maes, C.; Carmeliet, P.; Bouillon, R.; Carmeliet, G. Vitamin D receptor in chondrocytes promotes osteoclastogenesis and regulates FGF23 production in osteoblasts. J. Clin. Invest. 2006, 116, 3150–9.

31. Yagishita, N.; Yamamoto, Y.; Yoshizawa, T.; Sekine, K.; Uematsu, Y.; Murayama, H.; Nagai, Y.; Krezel, W.; Chambon, P.; Matsumoto, T.; et al. Aberrant growth plate development in VDR/RXR gamma double null mutant mice. Endocrinology 2001, 142, 5332–41.

32. Jiménez, M.J.; Balbín, M.; Alvarez, J.; Komori, T.; Bianco, P.; Holmbeck, K.; Birkedal-Hansen, H.; López, J.M.; López-Otín, C. A regulatory cascade involving retinoic acid, Cbfa1, and matrix metalloproteinases is coupled to the development of a process of perichondrial invasion and osteogenic differentiation during bone formation. J. Cell Biol. 2001, 155, 1333–1344.

33. Menéndez-Gutiérrez, M.P.; Ricote, M. The multi-faceted role of retinoid X receptor in bone remodeling. Cell. Mol. Life Sci. 2017, 74, 2135–2149.

